# Classic genome-wide association methods are unlikely to identify causal variants in strongly clonal microbial populations

**DOI:** 10.1101/2021.06.30.450606

**Authors:** Peter E. Chen, B. Jesse Shapiro

## Abstract

Since the advent of genome-wide association studies (GWAS) in human genomes, an increasing sophistication of methods has been developed for more robust association detection. Currently, the backbone of human GWAS approaches is allele-counting-based methods where the signal of association is derived from alleles that are identical-by-state. Borrowing this approach from human GWAS, allele-counting-based methods have been popularized in microbial GWAS, notably the generalized linear model using either dimension reduction for fixed covariates and/or a genetic relationship matrix as a random effect in a mixed model to control for population stratification. In this work, we show how the effects of linkage disequilibrium (LD) can potentially obscure true-positive genotype-phenotype associations (i.e., genetic variants causally associated with the phenotype of interest) and also lead to unacceptably high rates of false-positive associations when applying these classical approaches to GWAS in weakly recombining microbial genomes. We developed a GWAS method called POUTINE (https://github.com/Peter-Two-Point-O/POUTINE), which relies on homoplastic mutation to both clarify the source of putative causal variants and reduce likely false-positive associations compared to traditional allele counting methods. Using datasets of *M. tuberculosis* genomes and antibiotic-resistance phenotypes, we show that LD can in fact render all association signals from allele counting methods to be fully indistinguishable from hundreds to thousands of sites scattered across an entire genome. These classic GWAS methods thus fail to pinpoint likely causal genotype-phenotype associations and separate them from background noise, even after applying methods to correct for population structure. We therefore urge caution when utilizing classical approaches, particularly in populations that are strongly clonal.

## Introduction

To date, human genome-wide association studies (GWAS) have revealed meaningful evidence both about genetic architectures and the underlying pathways involved in human disease and other heritable traits. Microbial GWAS, while gaining popularity, is still in its infancy compared to human GWAS. Borrowing knowledge and methodology from human studies, microbial association studies are prone to similar pitfalls but also present new challenges due to the distinct and diverse population genetics of microbes [1]. Notably, population stratification and linkage disequilibrium (LD) present substantial impediments to identifying potential causal genotype-phenotype associations in strongly clonal populations. The first obstacle is a similar confounder seen in human GWAS, though microbial populations often exhibit a much higher magnitude of stratification owing to clonal descent. The second obstacle is the focus of this paper, and perhaps the greater obstacle of the two. Crucially, microbial populations exhibit both strong and long-range LD due to recombination mostly via relatively short gene conversion events rather than the process of crossing over, leaving large and potentially distant regions of the genome linked in a clonal frame [2].

The human haplotype map [3–5] exploited the block-like LD created by crossing over (i.e., recombination during meiosis in which homologous chromosomes exchange segments) to provide a shortcut where a relatively small subset of genome-wide markers within haplotype blocks could ‘tag’ other markers in LD as a proxy. The non-block-like structure of clonal frames seen in many microbial populations presents a situation that is the converse of that seen in the human haplotype map; where blocks allowed a shortcut to genome-wide coverage, clonal frames resemble one large haplotype block covering the length of entire microbial genomes, thus obscuring potential causal sites that are not distinguishable from the rest of the frame. This scenario is perhaps analogous to the fine-mapping problem where an attempt is made to clarify the source of the putative causal signal within an LD region [6], except in highly clonal populations the region to fine-map is the entire clonal frame. Population genetic theory informs us that, in genomes exhibiting strong LD, only sites under convergent selection, which experience homoplasic mutations in independent lineages and thus not in LD with other sites, are likely to be distinguishable from other sites [1]. Yet, much of the recent literature on microbial GWAS tends to report results without explicit mention of the LD profiles of the top hits, making it difficult to assess which mutations or genes are more likely to be driving the association and how many others are likely associated by LD.

Currently, GWAS methods can be broadly broken down into two general approaches based upon the source of their primary association signal: allele counting and homoplasy counting [1]. Regardless of the association model used, all allele counting methods derive their signal from alleles that are identical-by-state, whereas homoplasy counting methods derive their signal strictly from alleles that are identical-by-state but not identical-by-descent, often called homoplasic, convergent, or parallel mutations that arise repeatedly and independently on different genetic backgrounds. To date, most microbial GWAS in the literature have primarily relied on the classical allele counting methods invented for human studies, notably generalized linear regression using either dimension reduction for fixed covariates and/or a genetic relationship matrix as a random effect in a mixed model to control for population stratification [7–9].

Here we build off of our earlier homoplasy counting association method [10] and describe a next-generation GWAS method, which we call POUTINE. We apply POUTINE to two datasets of *M. tuberculosis* genomes and antibiotic-resistance phenotypes and find that it identifies known resistance mutations while minimizing likely false-positive associations. Utilizing our new tool, we further explore a major question concerning the state of classic allele counting and its use for strongly clonal microbial populations: Do allele counting methods identify signals outside of convergent sites, and are these likely true or false-positive associations?

## Results

### A new homoplasy counter

We begin by describing key aspects of POUTINE, a GWAS method based on homoplasy counting, with specific details elaborated in the Methods. Homoplasies at each nucleotide position (site) are identified by finding all identical alleles that do not share a most recent common ancestor. Using only homoplasic mutations offers solutions to two of the most substantial obstacles in microbial GWAS. First, homoplasic mutations having arisen on independent genetic backgrounds are not linked to other sites in the genome. This feature allows homoplasies to naturally bypass the LD problem because convergent sites are by definition unlinked from the clonal frame and thus provide truly independent association signals. Second, there is no need to further correct for the confounding effect of population stratification, which arises due to genetic ancestry. By definition, homoplasic mutations are not identical-by-descent, and thus do not contribute to spurious associations caused by subpopulations where cases are on average more genetically related with each other than controls. Avoiding population stratification correction preserves statistical power that is otherwise potentially lost due to this correction.

Our goal with POUTINE was to develop a homoplasy-based GWAS method that is both robust and user-friendly. Currently, POUTINE input phenotypes are strictly discrete and binary. Different from the original phyC method [10], ancestrally reconstructed phenotypic states are avoided due to the noise they may potentially introduce to the association signal. With the growing scale of genome sequencing, larger sample sizes can compensate for excluding ancestral genotypes and phenotypes from the homoplasy counts. Genotypes are strictly from the core genome and only biallelic single nucleotide variants (SNVs) are currently tested for associations with a discrete phenotype using a binomial test. The background mutation rate across the genome can potentially vary, thus allowing some sites to have a higher expected level of homoplasies. The binomial test incorporates the total number of observed homoplasies at each site, thus accounting for any varying background mutation rates. Critical to homoplasy counting is the robustness of the input tree topology as this directly determines which mutations are called homoplasic. The input ancestral phylogenetic reconstruction is optional; users may choose to precompute an ancestral reconstruction or opt for POUTINE to compute one based upon the topology of the input tree.

Homoplasic mutations are often taken as hallmarks of positive selection. The intuition is that adaptive mutations under positive selection will appear repeatedly in independent lineages experiencing the same selective pressure [11,12]. However, some baseline level of selectively neutral homoplasy is expected, and in the context of GWAS, some of these homoplasies will be spuriously associated with the phenotype under study. In POUTINE, we therefore establish an empirical null distribution (with no association between phenotype and genotype) by permuting the phenotypes and leaving the genotypic structures completely intact. We assess sites likely to be under convergent selection using Westfall and Young’s max(T) resampling scheme [13]. This marks an additional improvement from the earlier phyC method. Principally, it produces a familywise error rate (FWER) that is far less conservative than methods that treat each hypothesis as being independent. This feature is particularly appropriate in strongly clonal populations where many regions are in complete LD and thus all sites in the region present as identical hypotheses. In addition, in the context of strongly clonal populations, the max(T) method is also more favorable than classic false discovery rate (FDR) based approaches. Methods to strongly control the FDR, such as Benjamini and Hochberg’s step-up procedure [14] and Storey’s q-value [15], were originally proposed under the assumption of independent tests. As such, these methods are unproven to strongly control the FDR in the face of pervasive dependence structures between sites. We note that there have been recent developments in relaxing this assumption to allow for increasingly arbitrary dependence structures [16–20].

As a further improvement upon phyC, POUTINE has been designed to handle much larger datasets. To scale to larger sample sizes in the tens of thousands of genomes, our implementation of POUTINE uses several optimizations including parallelizing the most time-consuming step, which is resampling. To further decrease runtime, the user may set a minimum homoplasy count to ignore sites with low counts that are unlikely to be statistically significant. We find this option to be helpful as many sites potentially have only one or two homoplasic mutations.

### Benchmarking POUTINE on a test dataset

To empirically test the sensitivity and specificity of POUTINE, we reanalyzed a ‘test’ dataset of 123 *M. tuberculosis* genomes previously analyzed with phyC to identify convergent mutations associated with a broad resistance phenotype (defined as resistance to any anti-TB drug by conventional drug susceptibility testing) [10], and compared our new results to this reference set. Table S1 shows the six genome-wide significant hits from the reanalysis. All four genes previously identified in the literature as causal genes for drug resistance (*rpoB, embB, rpsL, rrs*) were re-identified as top hits, except for *rpsL*. The lack of signal for *rpsL* could be due to the relatively small sample size of 123 genomes and only four homoplasic mutations at this site, not including mutations in ancestrally reconstructed internal branches. By including internal branches, the original phyC may have gained power to identify associations in *rpsL*, but at the likely risk of additional false-positive associations. In particular, phyC identified associations in 16 PE/PPE genes in this dataset, which were reasoned to be false-positive associations [10], while the POUTINE reanalysis did not identify a single association in a PE/PPE gene. The family of PE/PPE genes in *M. tuberculosis* is known to be problematic for both sequencing and alignment due to their similarity and repetitive nature; thus they are prone to GWAS false positives [10]. Three additional genes were identified by POUTINE that were not seen in the previous analysis (Rv0853c, Rv0587, Rv1639c). An examination of the literature reveals that each of these three genes has been implicated in drug resistance in *M. tuberculosis* [21–23]. Overall, this real-world example suggests that POUTINE is similarly sensitive compared to phyC (detecting likely true positives), and also more specific (reducing likely false positives in PE/PPE genes).

### An LD perspective on allele and homoplasy counting signals

To compare the LD profiles of convergent vs. non-convergent GWAS hits, we analyzed a second ‘discovery’ dataset of 1330 *M. tuberculosis* genomes and an isoniazid drug-resistance phenotype [24]. To identify non-convergent sites, we used a standard allele-counting GWAS approach, implemented in PLINK using logistic regression with principal components as covariates to control for population stratification (Methods).

Using POUTINE, we identified three SNVs significantly associated with isoniazid resistance after correcting for multiple hypothesis testing, and another three ‘secondary’ hits which did not survive multiple test correction (max(T)-corrected *P* = 0.058; Table 1). For illustrative purposes, we consider these secondary hits because all three sites are one homoplasic case count away from genome-wide statistical significance, suggesting that a larger sample size or combining more drug-resistant phenotypes into one broad resistant phenotype would show these sites to be significant. In addition, two of the three secondary hits (sites 1674048 and 1674481) are in the same region as the primary hit at site 1673425, suggesting that a set-based test (i.e., combining mutation counts in the same gene or region) would likely identify these two secondary hits as genome-wide significant.

**Table 1.**
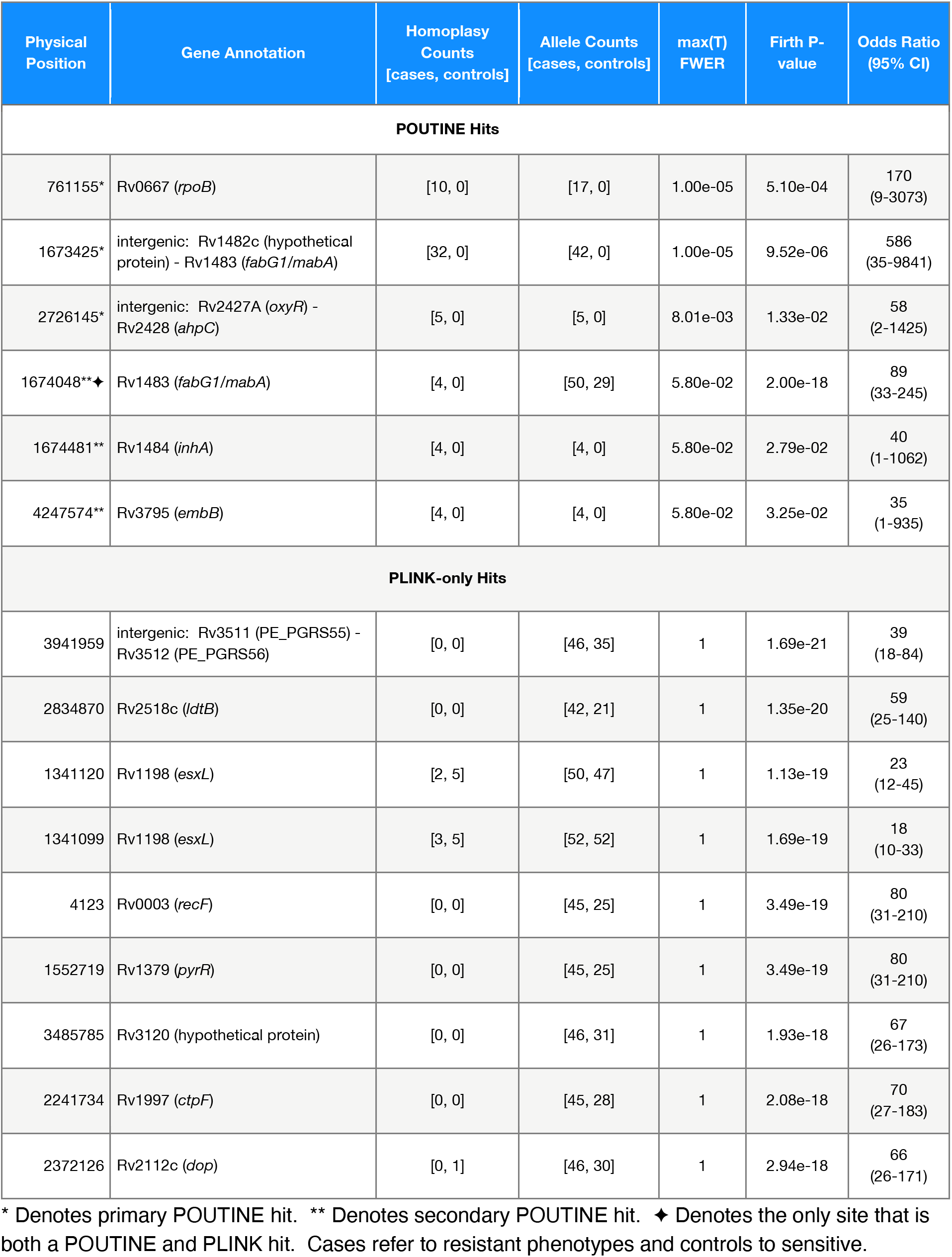
Discovery set top hits (POUTINE + PLINK-only hits).

In contrast to POUTINE, it proved difficult to distinguish GWAS hits from background noise using PLINK, even with standard corrections for population stratification. Due to pervasive LD, a genome-wide significance threshold cannot be relied upon to identify a subset of plausible hypotheses because of the overwhelming numbers of false positives that LD drags below this threshold. Consider a Bonferroni-corrected *P*-value cutoff of 0.01 (red line in Figure 1). Even using such a conservative threshold would still include too many false-positive associations and would leave the investigator with 288 genome-wide significant PLINK hits – a daunting number to consider for experimental follow-up. Manhattan plots of strongly clonal populations often feature groupings of sites in strong LD that we refer to as ‘LD frames’, visible as horizontal lines of sites with near-identical *P*-values (Figure 1). We refer to these plots as Montreal plots to reflect the low, horizontal skyline in contrast to the vertical skyscrapers of Manhattan.

**Figure 1.**
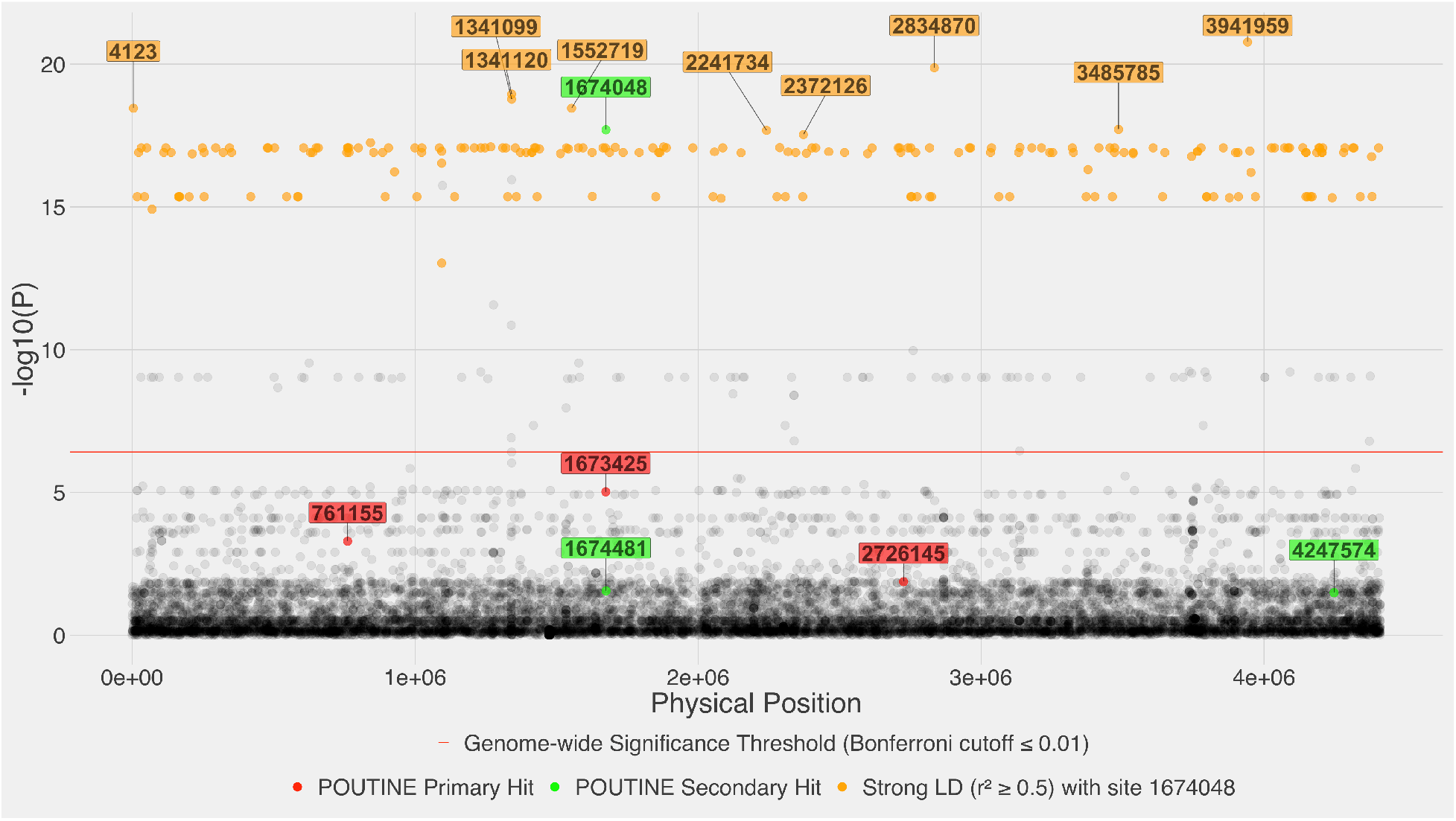
Montreal plot of the discovery set with correction for population stratification. *P*-values along the y-axis were calculated using PLINK’s logistic regression with the Firth correction. Population stratification was corrected using the first four principal components. The x-axis shows the nucleotide positions along the *M. tuberculosis* genome.

For our purpose of examining LD profiles of allele counting hits, we arbitrarily chose the 10 sites visually distinguishable above the top LD frame as the top PLINK hits (Figure 1). Of these 10, only site 1674048 overlaps with a secondary POUTINE hit (Table 1). The careful reader will note that the top PLINK hit by raw allele counts (42 cases to 0 controls in Table 1) should be site 1673425 but this is not the case in Figure 1. Due to the phenomenon of complete statistical separation at this site (caused by allele counts showing all cases and zero controls) as well as a small sample size at the minor allele, even the Firth correction used does not sufficiently improve the accuracy of the significance estimate (note in Table 1 the wide confidence intervals around the estimated effect size). However, this problem is avoided when using an exact test such as Fisher’s exact test, for which we do see site 1673425 as the top hit (Figure S2). For the purposes of the further analyses below, this detail does not affect our general conclusions.

To further investigate the effects of linkage on GWAS hits, we compared how PLINK-only hits were linked to other sites in the genome to the six POUTINE hits. To focus on relatively strong LD, we plot sites across the genome that are linked to a top GWAS hit with a threshold of r^2^ >= 0.5 (Figure 2). The complete distribution of r^2^ values is shown in Supplementary Figure S1. Contrasting the two sets of LD profiles shows that the homoplasy-counting-based POUTINE signals are predominately free of strong linkage from each other and from the rest of the genome. Conversely, the allele-counting hits show strong linkage to sites throughout the genome. The only POUTINE hit that shows a similar LD pattern to the set of PLINK-only hits is position 1674048. This is the only POUTINE hit where the mutations are predominantly non-homoplasic; there are 79 minor alleles at this site, only four of which are homoplasies (Table 1). The other five POUTINE hits are at sites that consist of entirely or predominantly homoplasies. These five hits suggest that sites composed of predominantly homoplasic mutations can be considered the sole source of an association signal, i.e., there are no other sites in strong LD that can be driving this signal, or hitchhiking along with a causal association. In contrast, the associations at site 1674048 and the set of nine PLINK-only hits cannot be disentangled from linked mutations across the genome (Figure 2). Because these hits show strong linkage to each other and to many sites across the entire genome, it is unclear how many and which of these sites are driving the association signal.

**Figure 2.**
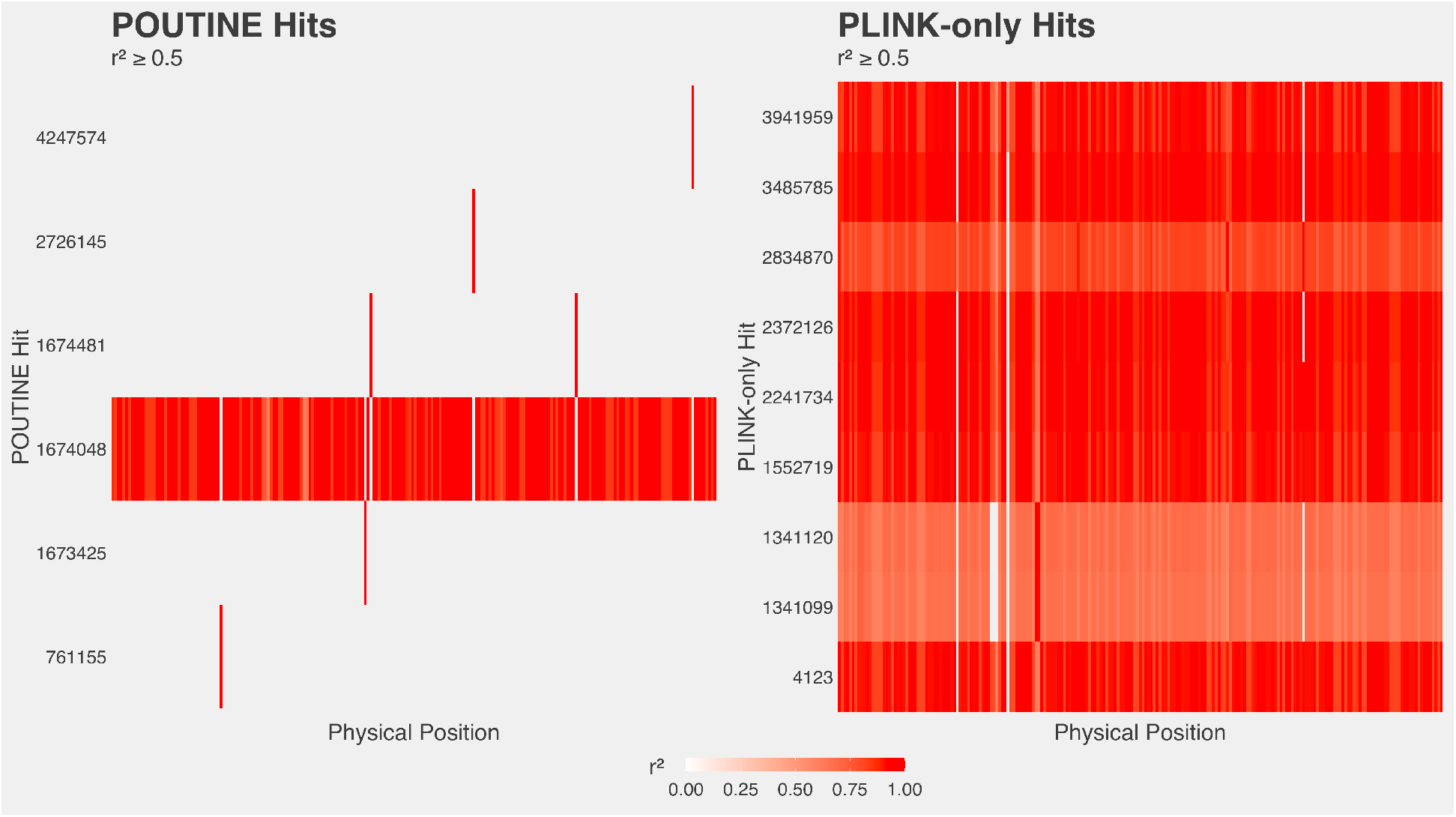
Genome-wide LD (r^2^ >= 0.5) of top hits (POUTINE + PLINK-only hits).

To determine what might be driving the POUTINE hit at site 1674048, we identified all the sites to which it is strongly linked (r^2^ >= 0.5). Strikingly, all sites in strong LD with site 1674048 (orange points in Figure 1) include the entire top LD frame and all nine PLINK-only hits. Among these 10 potential hits, site 1674048 is the only one with a known causal mutation (a silent mutation that confers isoniazid resistance) in the literature [25]. It, therefore, seems likely that this site is driving the association signal, with the other sites being associated due to linkage.

What would the association results look like if one relied only on classical allele counting methods (i.e., all homoplasy information is removed from the above analyses)? First, one would mistakenly identify 9 of the 10 top hits as plausible candidates of association to isoniazid resistance, with site 1674048 being the only known causal mutation. Even upon further examination of LD for all 10 sites, one is left with no meaningful signal because each top hit is strongly linked to sites across the entire genome; the likely causal site 1674048 does not stand out in any identifiable way. Second, one would miss all five other POUTINE hits (sites 761115, 1673425, 1674481, 2726145, 4247574). Since these five sites are predominantly homoplasic, it is possible that with increased sample sizes allele counting would identify them, provided that the increase in sample size also increases the minor allele frequencies at these sites. Only then could further examination of LD reveal that these five sites are not strongly linked to one another nor to the rest of the genome, and thus be considered as plausible candidates.

### Population stratification corrections do not break up LD

As an alternative to homoplasy counting, allele-counting GWAS methods are typically corrected in an attempt to remove the confounding effect of population stratification. Regardless of the population stratification control used, these corrections effectively reweight statistical significance at each affected site and thus only shifts the site up or down along the y-axis of a Manhattan plot. Crucially, these corrections say nothing about the correlational structures between sites. Consider how the Manhattan plot changes when one removes the stratification control; here we simply removed all principal components used in the regression model (Figure 3). The nine PLINK-only hits identified with stratification correction (Figure 1; Table 1) are no longer discernible from the top LD frame, leaving only site 1674048 as the lone top hit which rises above the top LD frame in the absence of correction (Figure 3). Therefore, stratification correction of an allele counting GWAS identifies eight additional hits compared to an uncorrected approach, none of which overlap with homoplasy counting hits. Without LD information, one is susceptible to being misled into thinking that these allele counting hits are distinguishable from the top LD frame, when in fact their LD profiles say otherwise (Figure 2). In summary, stratification correction alone cannot bypass the confounding effects of LD.

**Figure 3.**
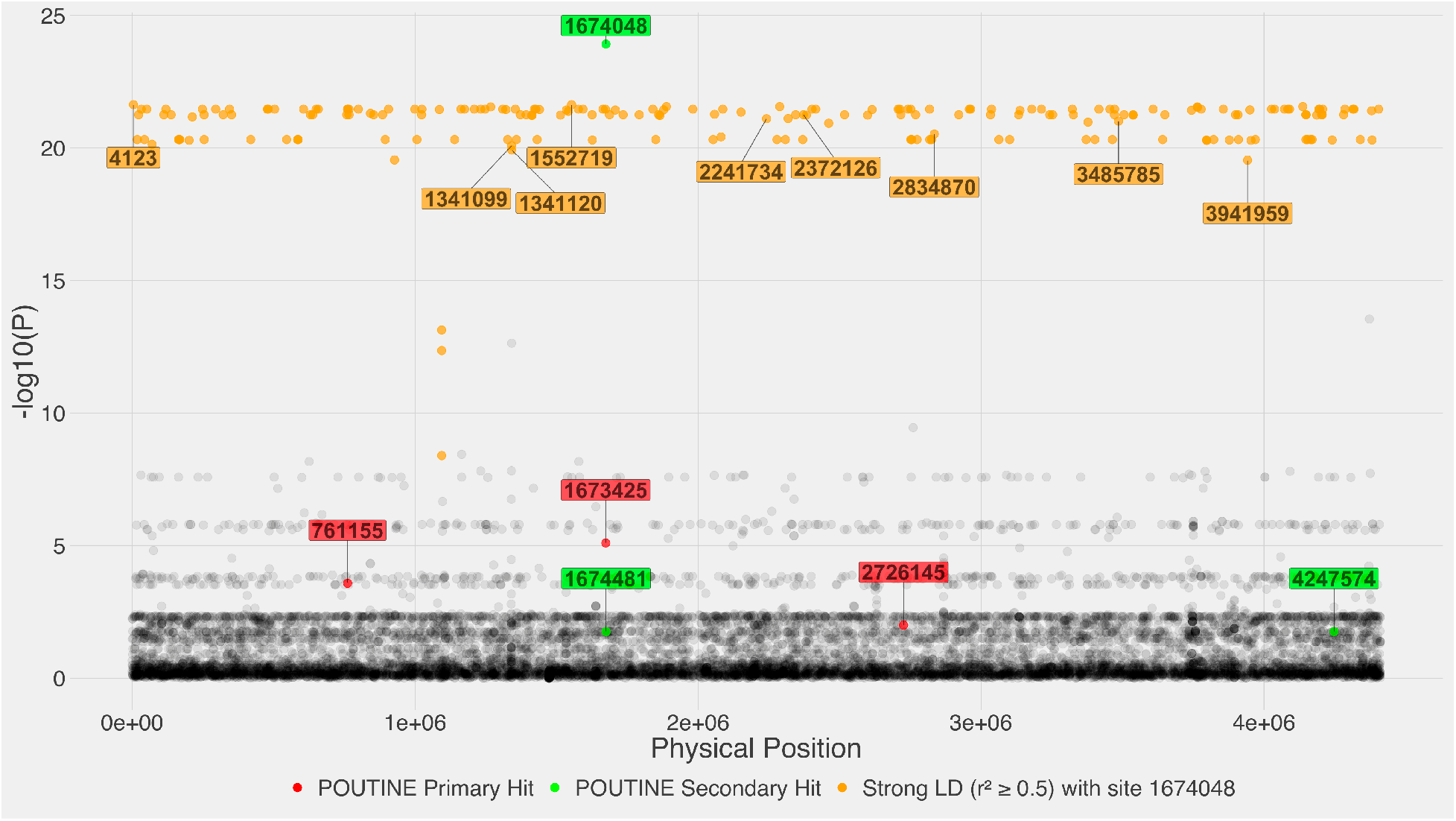
Montreal plot of the discovery set without correction for population stratification. The analysis is identical to Figure 1, except no population stratification correction was done.

## Discussion

### The state of allele counting methods for strongly clonal populations

To demonstrate the impediment LD presents to classical GWAS methods, we chose *M. tuberculosis*, a clonal species that features strong and long-range LD across the entire genome. We sought to answer a critical question regarding the utility of homoplasy- and allele-counting GWAS approaches: Do allele counting methods meaningfully identify signals outside of sites evolving under convergent evolution? The answer to this question is likely no. Although allele counting methods may produce strong GWAS signals for causal variants that are not convergent, those signals are indistinguishable from other similarly strong signals in linked sites. Crucially, the vast majority of these linked sites – which can easily number in the hundreds or thousands – are likely false positives. The LD frames that represent groupings of strongly linked sites feature prominently in Manhattan plots of strongly clonal microbial populations. In such populations, Manhattan plots are more appropriately called Montreal plots because they more closely resemble the relatively flat skyline of a city like Montreal (where regulations prevent buildings taller than its namesake Mount Royal, with an elevation of only 233 m) rather than the skyscrapers punctuating the Manhattan skyline.

### Is LD friend or foe?

The implications of these findings can be extended to other clonal populations, and likely to many other populations across the gamut of microbial recombination rates. It is unclear if there exist microbial populations with a sufficiently high rate of recombination to exhibit similar block-like LD structures seen in eukaryotic genomes. If so, these populations would be more amenable to allele counting methods. We note that there do exist recombination hotspots in bacterial species, such as *E. coli* [26], *C. jejuni* [27], and others [28]. These distinct regions are relatively unlinked from the rest of the genome, and as such can potentially provide a cleaner signal of association. In one such notable example, Chewapreecha *et al*. analyzed a highly recombining *S. pneumoniae* population to identify six common recombination hotspots [29]. A follow-up GWAS applying an allele-counting approach (specifically, the Cochran-Mantel-Haenszel test using population clusters identified with BAPS [30]) on beta-lactam antibiotic resistance in this population identified plausible association signals in genes encoding penicillin-binding proteins and involved in peptidoglycan synthesis – which tended to reside within one of the six common recombination hotspots [31]. It remains to be seen if allele-counting GWAS approaches can identify hits outside of such hotspots.

A viral population of current substantial interest is SARS-CoV-2. This population offers a timely example of the ramifications of our findings. This population is strongly clonal with evidence showing that there has been little realized recombination, although the potential for recombination is present [32]. Intriguingly, many of the variants of concern (VOCs) identified thus far harbor mutations (e.g., E484K in the spike gene) that are thought to be under convergent selection [33]. This highlights that even potentially recombining populations may be effectively clonal at the early stages of an outbreak (or pandemic), at which time homoplasy-counting methods are likely to be much more effective than allele-counting to identify genotype-phenotype associations.

### Limitations of POUTINE and future directions

Homoplasy-based methods such as POUTINE provide a promising lifeline for tackling association studies in microbial populations. Its major limitation is that if the causal variants to discover are not convergent mutations, then there simply will be no signal to discover. It is unclear at this time what proportion of causal variants are sculpted by convergent evolution. In addition, it is also unclear how much genetic heterogeneity underlies the genetic architecture of many microbial traits. Both locus and allelic heterogeneity can dilute the association signal, requiring higher sample sizes to recapitulate any signal. To address allelic heterogeneity, we plan on adding a set-based test to aggregate individual variants into localized regions to boost the signal. Currently, our initial implementation assays biallelic SNVs inside the core genome. As such, POUTINE excludes sources of variation including tri/quad-allelic sites, small indels, and the accessory genome. In the future, it would be straightforward to include tri/quad-allelic sites using a multinomial test instead of the binomial. We note that it is currently possible to recode loci in the accessory genome as present or absent among the population and run POUTINE as if these recoded loci were core SNVs. When taking this approach, one should proceed with caution because the recombination dynamics of the accessory genome may differ from the core. Lastly, a further limitation of homoplasy-based methods can be their inability to identify hemiplasies from homoplasies. A hemiplasy is a form of incomplete lineage sorting that can mask as a homoplasy [34,35]. If hemiplasies were mistakenly identified as homoplasies, it would no longer be appropriate to consider a homoplasy-based hit to be free from linkage to other sites. As such, sites composed of hemiplasies would present the same problem to homoplasy counting methods as that seen in non-convergent sites for allele counting methods.

This work highlights the necessity to both examine and report the LD profiles of top association signals. However, the importance of examining LD should not steer the reader away from utilizing prior knowledge of their phenotype of interest, as such knowledge can play a clarifying role in narrowing down hypotheses. However, lesser understood phenotypes serve as prime targets for the agnostic view of GWAS to reveal underlying mutations in genes and other loci we know nothing or little about.

Because many causal variants may be hiding in non-convergent sites, it is critical that we understand if allele counting methods can provide meaningful association signals in the face of pervasive LD observed across microbial populations. For strongly clonal populations that are not amenable to allele counting approaches, we must improve upon these classical methods, and if these methods prove intractable to LD, we must open a new line of inquiry perhaps beyond homoplasy-based solutions in hopes of capturing non-convergent causal variants. Until such time, we urge caution when using classical GWAS methods to tackle microbial populations, particularly those with little measurable recombination.

## Methods

### Sample collection, genotyping, and phenotyping

The reference set of 123 *Mycobacterium tuberculosis* genomes comprises 14 major phylogenetic clusters from different micro-epidemics and 23 geographically diverse drug-sensitive isolates. Of the 123 isolates, 47 were resistant to one or more antibiotics. Isolate selection, sequencing, variant calling, and phenotyping are described in detail in [10].

The discovery set of 1330 *M. tuberculosis* genomes (GenBank BioProject accession: PRJNA413593) includes isolates collected by the British Columbia Public Health Laboratory of the British Columbia Centre for Disease Control. Isolate collection and phenotypic drug susceptibility testing are detailed in [24]. Samples were sequenced on the Illumina HiSeq2500 platform at the Michael Smith British Columbia Genome Sciences Centre using 125-bp paired-end reads [36]. All reads were quality checked using FastQC [37], trimmed with Trimmomatic [38], and mapped against the H37Rv reference genome (GenBank Reference Sequence accession: NC000962.2) using BWA-MEM [39]. All variants were called using GATK [40] requiring a Phred quality score > 20 and read depth > 5. Variants were further filtered out if a site had a missing call rate > 10% of samples, and only biallelic SNVs were kept.

### Allele counting

Both PLINK 1.9 and 2.0 [41] were used for both preprocessing and association testing described below. All PLINK analyses were done with PLINK version 1.90b6.21 64-bit (19 Oct 2020) except for firth regression which used PLINK version 2.00a3 AVX2 (28 Mar 2021). For all runs, --chr-set −1 was used to designate a single haploid genome.

### Population substructures

To capture population substructures, LD pruning was done in two ways: 1) whole-genome pruning using one window including all markers, and using the PLINK option --indep-pairwise with varying r^2^ thresholds of 0.50, 0.90, 0.99; 2) local pruning using non-overlapping windows of 1000 markers at a time, and using --indep-pairwise with an r^2^ threshold of 0.99. Principal component analyses were run on the above LD-pruned sets and once without any LD pruning using --pca 20 header var-wts (Figure S3).

### Associating testing

Fisher’s exact testing was run with a mid-p correction and 10^7^ permutations using PLINK options --assoc fisher-midp mperm=10000000, and filtering out sites with a minor allele count < 3, sites with a missing call rate > 0.10, and samples with a missing call rate > 0.10 using --mac 3, --geno 0.1, --mind 0.1, respectively.

Logistic regression with a Firth penalty was conducted using --glm firth cols=+a1count,+totallele,+a1countcc,+a1countcc,+totallelecc,+a1freq,+a1freqcc. Firth correction serves as a penalty during maximum likelihood estimation to avoid nonconvergence issues due to statistical separation [42].

To control for population stratification, --covar was used with the LD-pruned set, described above, which was calculated with non-overlapping windows of 1000 markers. The first four principal components were selected as fixed covariates using --covar-name PC1-PC4 (Figure S4). Sites were removed from Firth regression using --mac 4, --geno 0.1, and --mind 0.1. Multiple-testing correction reports were generated using --adjust. 95% confidence intervals around each odds ratio were reported using --ci 0.95.

### Linkage disequilibrium

r^2^ was calculated using PLINK options --r2, --ld-snps followed by sample ID names, --ld-window 69722 and --ld-window-kb 5000 to remove default settings to allow all genome-wide pairs of markers conditional on the specified sites in --ld-snps, and --ld-window-r2 0 to report all r^2^ values.

### Homoplasy counting

All steps except for the building of an input tree are implemented in POUTINE, the method introduced here.

### Input tree

Phylogenies were inferred using both FastTree [43] version 2.1.11 double precision (No SSE3) and raxml-ng [44] version 1.0.2. FastTree was run with -nt -gtr settings, and raxml-ng was run with --model GTR+G settings. To improve the inference of the tree topology, likely homoplasic regions (39 genes previously associated with drug resistance from [45]), as well as repetitive regions (e.g., PE/PPE and PGRS genes), were filtered out (273 genes; 10% of the genome; genes listed in [46]). Both tree topologies were effectively identical and did not affect the final homoplasy-based hits (Figure S5). Newick format parsing was done using the Coevolution library [47].

### Homoplasy identification

Ancestral genotypic reconstructions were done using TreeTime [48] version 0.7.6. The ancestral subcommand was used with the --gtr infer settings and used the default joint maximum likelihood method. To identify homoplasic mutations, ancestral mutations were mapped onto the input tree, and a homoplasic event was called if at least two identical mutations/alleles at the same segregating site occurred on independent genetic backgrounds, i.e., the two mutations do not share a most recent common ancestor.

### Association model

Associations of homoplasic mutations to the phenotype of interest is conducted using a binomial test using the binomial distribution defined as binomial(n, p, x), where n is the total number of homoplasic mutations observed at each site, p is the probability of success of each Bernoulli trial and is equal to the case to control ratio for all sites, and x is the count of homoplasic mutations in cases at each site. We assume that the mutation rate at a particular site is a proxy for the background rate of expected homoplasic mutations at that site. Considering n at each site helps address any possible variation in the background mutation rate across the genome, and also satisfies the exchangeability principle between sites to allow for resampling of permutations. The implementation of the binomial test is from the Apache Math Commons 3.6.1 library (https://commons.apache.org/proper/commons-math/).

### Significance assessment

*P*-values are derived using a resampling-based multiple-testing method by Westfall & Young [13]. This approach is sometimes referred to as max(T) for the maximum test statistic. max(T) is a single-step adjustment method:

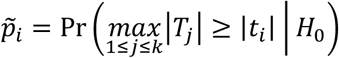

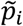 is the probability that the largest test statistic in the resampled data set, *T_j_*, is larger than the observed test statistic, *t_i_*, given that all null hypotheses, *H*_0_, are true. *k* is the number of tests.

Specifically, for each site a familywise error rate (FWER) is calculated as follows:

1. pointwise estimate: for each replicate, only phenotypes are permuted while genotypes and their dependency structures are left unmodified and completely intact. For each site, a resampled null distribution of no association is constructed from m replicates, where m is defined as the user-specified total number of replicates/permutations. The pointwise estimate is defined as (*r_point_* + 1)/(*m* + 1) [49], where *r_point_* is the number of replicates equal to or more extreme than the observed binomial test statistic.
2. familywise estimate: for each replicate, the maximum test statistic is saved, and a second null distribution is constructed using all m maximum test statistics across all replicates. The familywise estimate is defined as (*r_family_* + 1)/(*m* + 1), where *r_family_* is the total number of maximum test statistics equal to or more extreme than the observed binomial test statistic.

## Availability of POUTINE

Download and installation instructions for POUTINE are available at GitHub: https://github.com/Peter-Two-Point-O/POUTINE.

## Acknowledgments

We would like to thank Jennifer Gardy and Ben Sobkowiak for their assistance with the discovery set (GenBank BioProject accession: PRJNA413593). We further thank James Tanner, Len Taing, Arnaud N’Guessan, and Gavin Douglas for alpha testing our software and also for helpful discussions and valuable feedback.

## Supplementary material

**Table S1.** All six genome-wide significant POUTINE hits in the reference set of 123 *M. tuberculosis* genomes.

| Physical Position | Gene Annotation | Homoplasmy Counts [cases, controls] | max(T) FWER |
| --- | --- | --- | --- |
| 1473246* | Rvnr01 (rrs) | [9,0] | 8.89e-04 |
| 761155* | Rv0667 (rpoB DNA-directed RNA polymerase subunit beta) | [8,0] | 6.30e-03 |
| 949535** | Rv0853c (pdc alpha-keto-acid decarboxylase) | [8,0] | 6.30e-03 |
| 4247429* | Rv3795 (embB arabinosyltransferase B) | [7,0] | 2.90e-02 |
| 685461** | Rv0587 (yrbE2A hypothetical protein: ABC-type transporter Mla maintaining outer membrane lipid asymmetry, permease component MlaE [Cell wall/membrane/envelope biogenesis]) | [7,0] | 2.90e-02 |
| 1847919** | Rv1639c (hypothetical protein: Enterochelin esterase or related enzyme [Inorganic ion transport and metabolism]) | [8,1] | 4.23e-02 |

\* Denotes an overlapping hit with phyC. \*\* Denotes a new POUTINE hit not identified by phyC.

**Figure S1.**
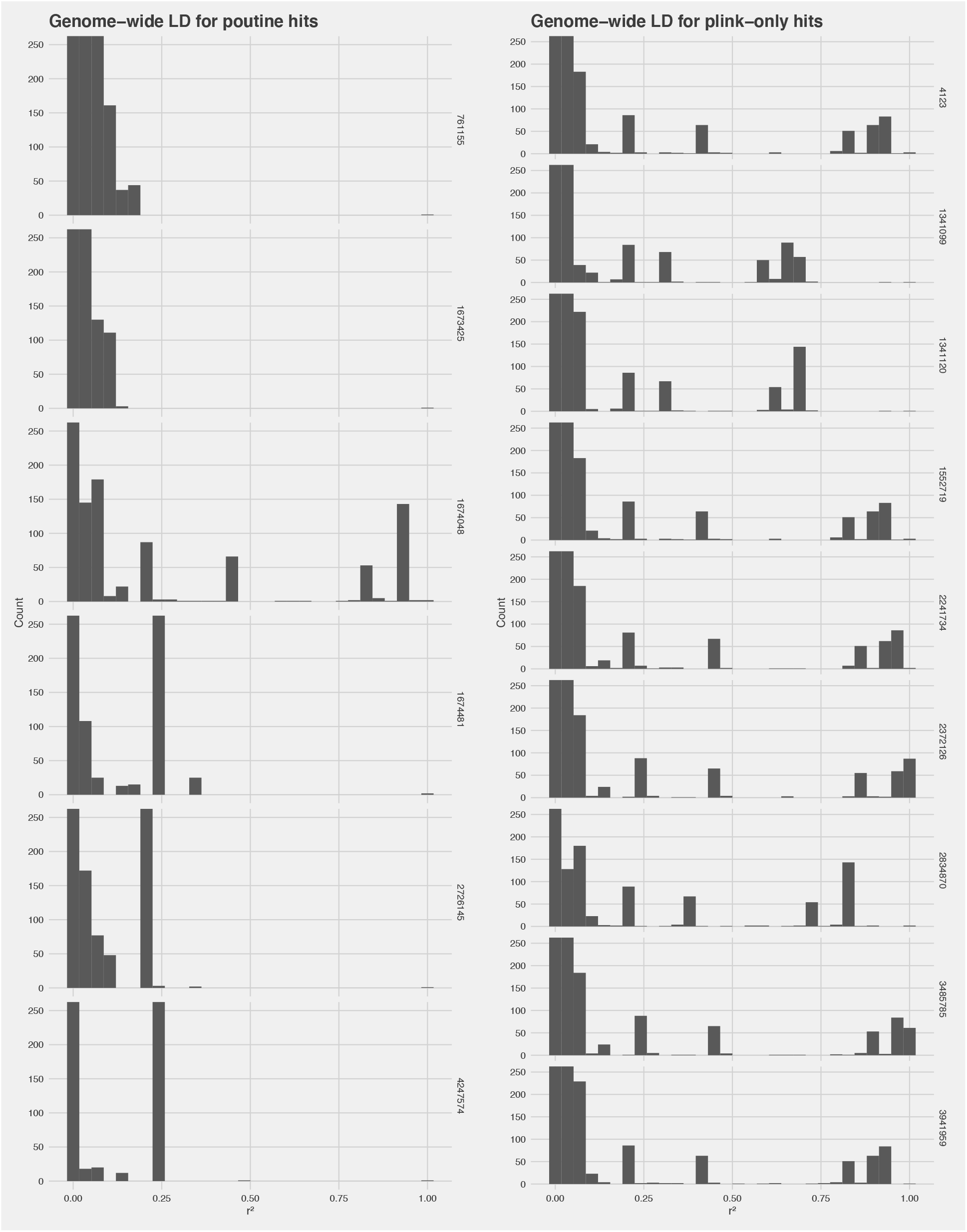
Distribution of genome-wide *r^2^* values for the set of POUTINE hits vs the set of PLINK-only hits.

**Figure S2.**
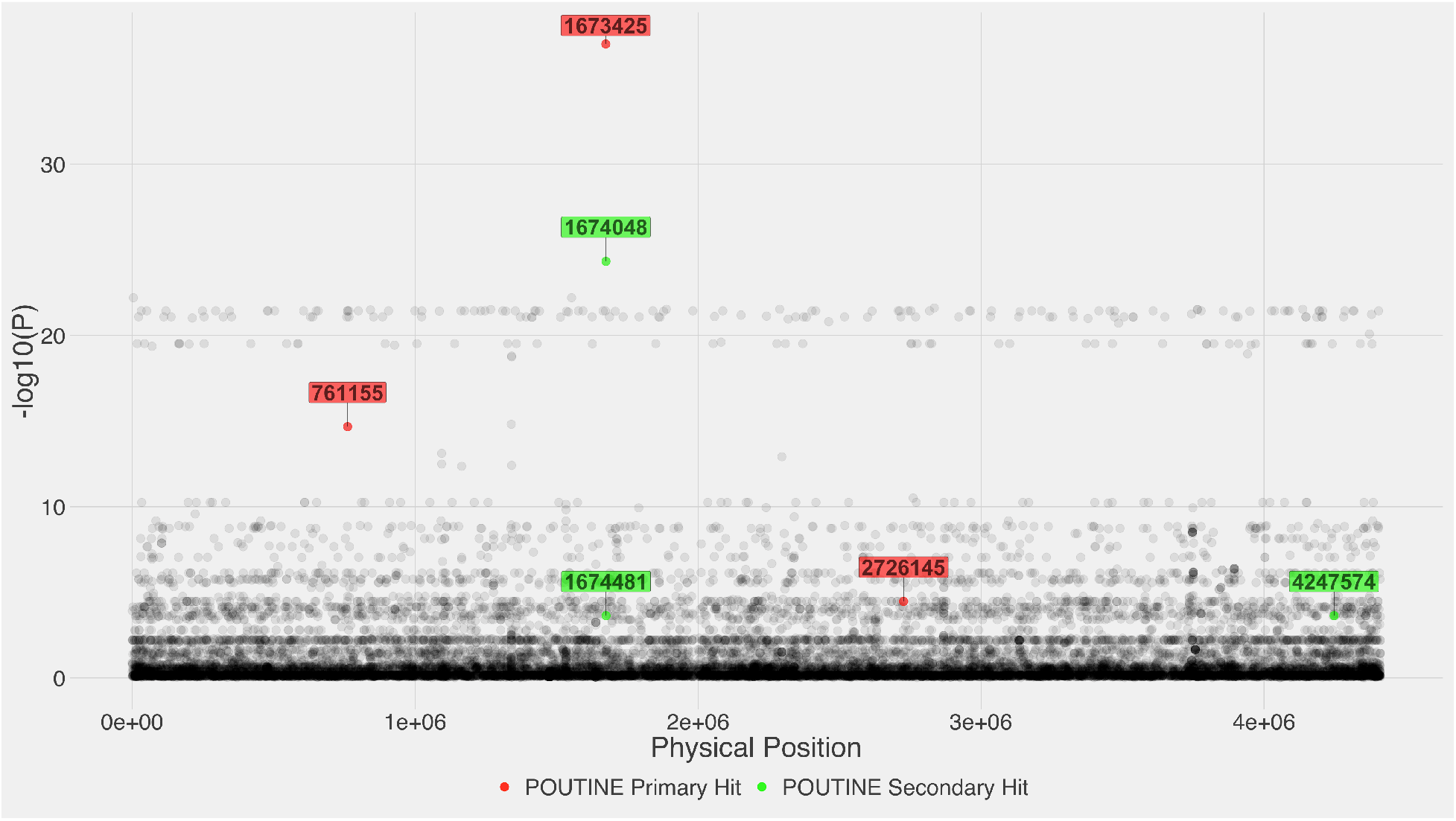
Montreal plot of the discovery set. *P*-values along the y-axis were calculated using PLINK’s Fisher’s exact test with the mid-p correction.

**Figure S3.**
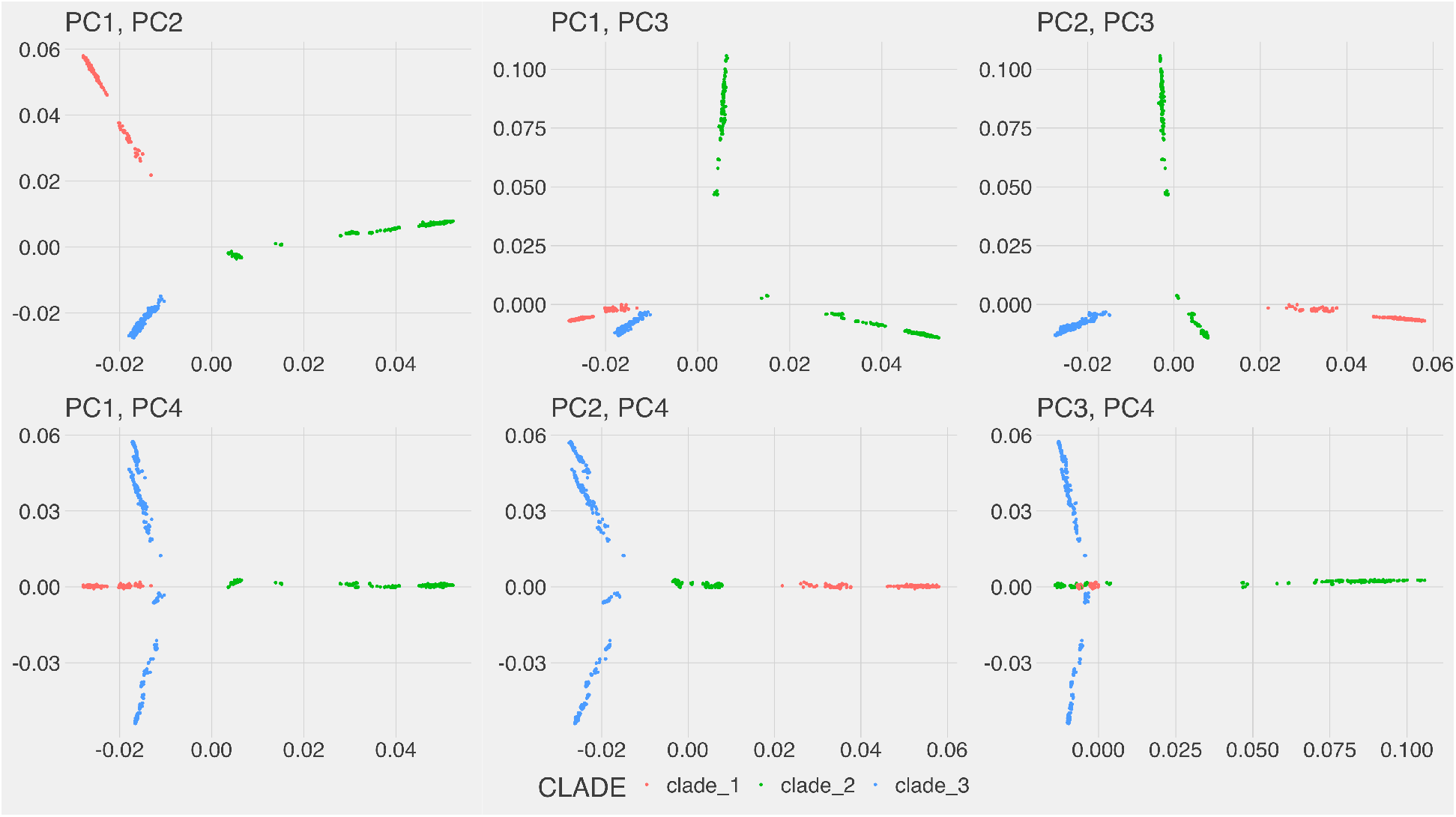
Score plots of the first four principal components derived from the LD pruned set using r^2^ > 0.99 and non-overlapping windows of 1000 sites. The three colors (red, green, and blue) highlight three broad subpopulations as inferred from the phylogeny of the discovery set (Figure S5).

**Figure S4.**
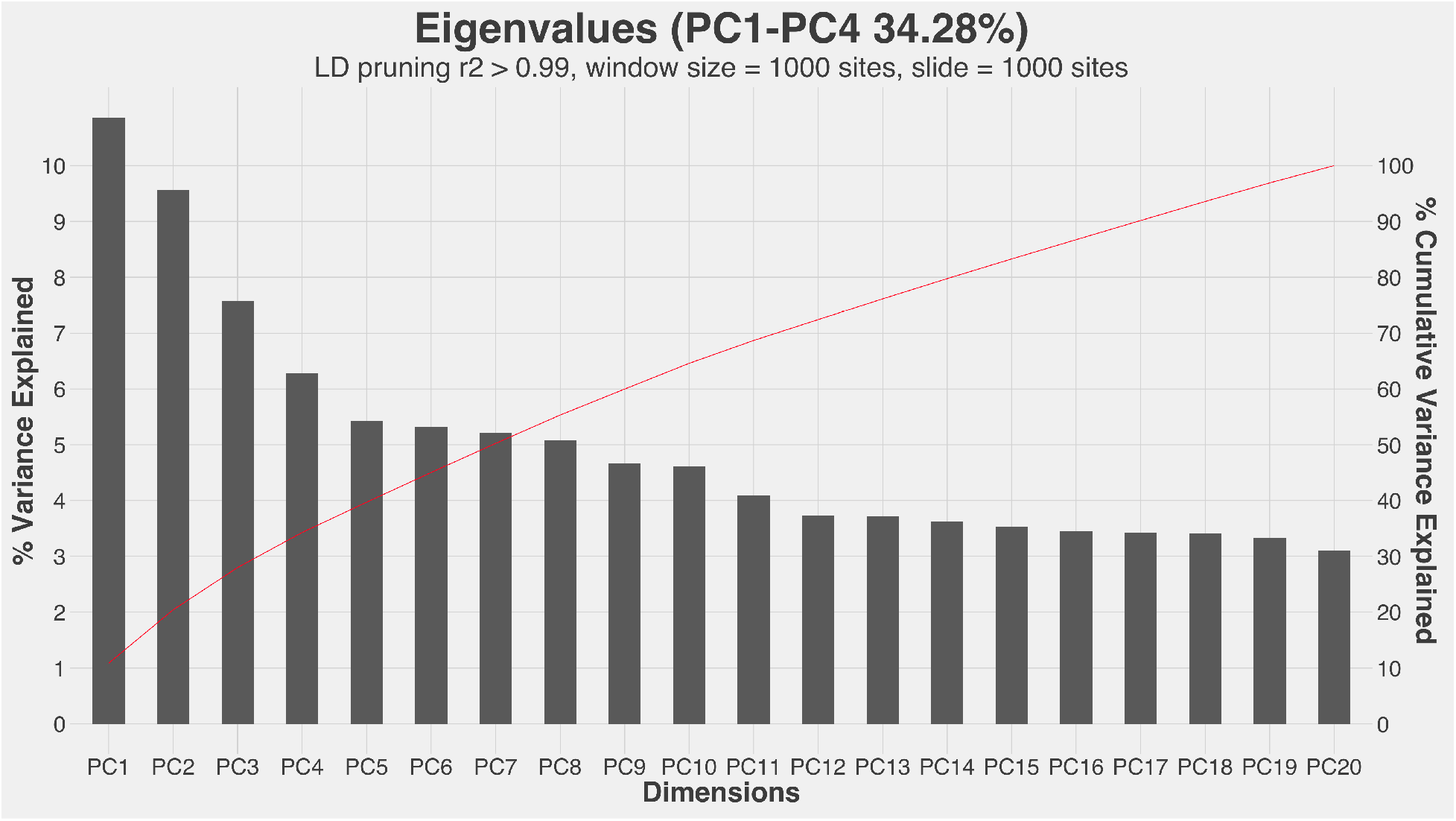
Scree plot of the first 20 principal components derived from the LD pruned set using r^2^ > 0.99 and non-overlapping windows of 1000 sites.

**Figure S5.**
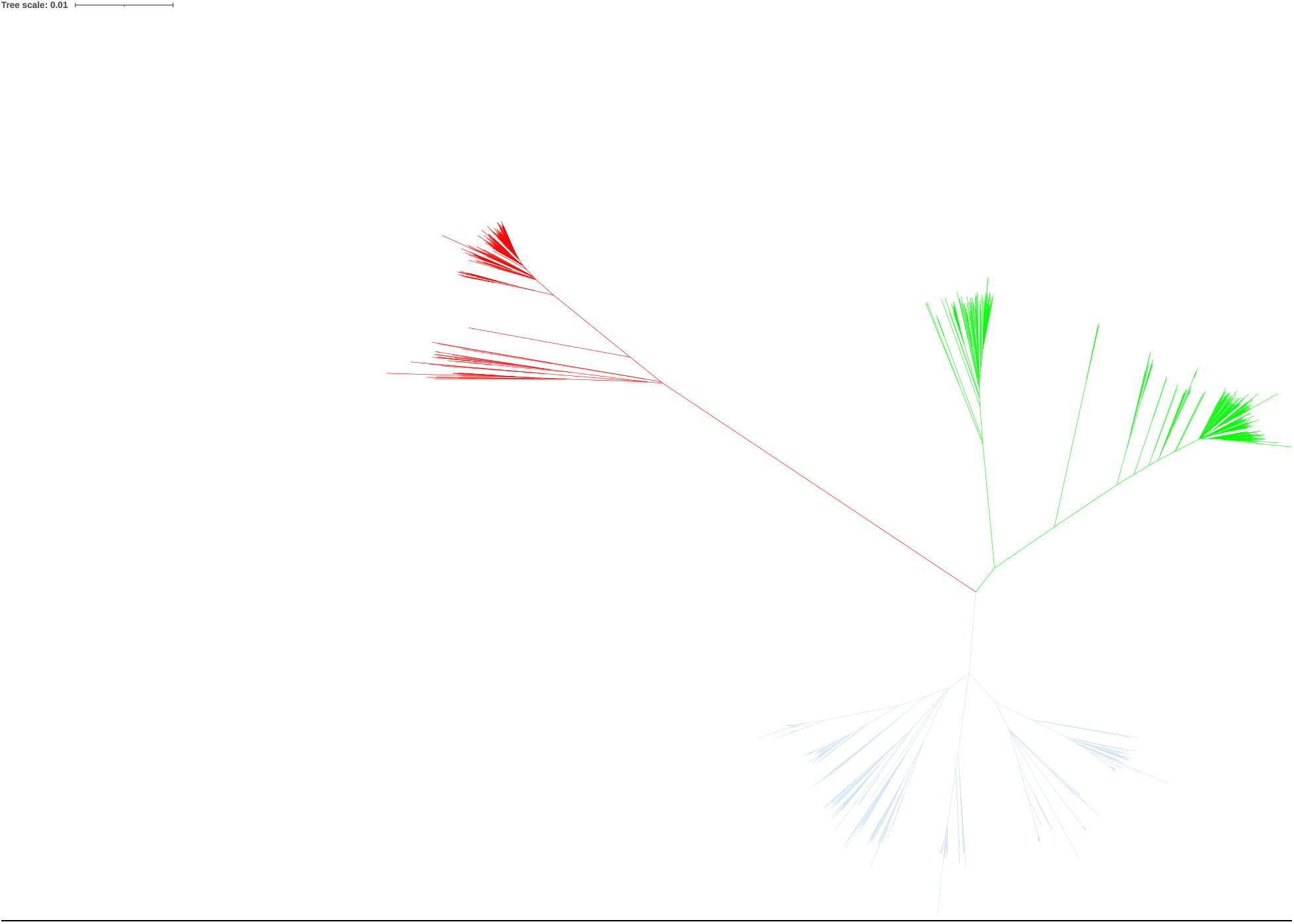
Phylogeny inferred using FastTree (double precision version) of the discovery set of 1330 *M. tuberculosis* genomes. The three colors (red, green, and blue) highlight three broad subpopulations.

